# Oviduct epithelial cells constitute two developmentally distinct lineages that are spatially separated along the distal-proximal axis

**DOI:** 10.1101/2020.08.21.261016

**Authors:** Matthew J Ford, Keerthana Harwalkar, Alain S Pacis, Helen Maunsell, Yu Chang Wang, Dunarel Badescu, Katie Teng, Nobuko Yamanaka, Maxime Bouchard, Jiannis Ragoussis, Yojiro Yamanaka

## Abstract

Owing to technical advances in single cell biology, the appreciation of cellular heterogeneity has increased, which has aided our understanding of organ function, homeostasis and disease progression. The oviduct (also known as the fallopian tube in humans) is the distal-most portion of the female reproductive tract. It is essential for reproduction and the proposed origin of high grade serous ovarian carcinoma (HGSOC). In mammals, the oviduct is morphologically segmented along the ovary-uterus axis into four evolutionally conserved regions. It is unknown however if there is a diversification of epithelial cell characteristics between these regions. In this study, we identified transcriptionally distinct populations of secretory and multiciliated cells restricted to the distal and proximal regions of the oviduct. We demonstrated that these distal and proximal populations are distinct lineages specified early in Müllerian duct development and are maintained separately. These results aid our understanding of epithelial development, homeostasis and initiation of disease from the oviduct.

## Introduction

The oviduct is the conduit connecting the ovary to the uterus and plays an essential role in promoting fertilization and preimplantation embryonic development (Shuai Li & Winuthayanon, 2017). With the advent of *in vitro* fertilisation (IVF) technologies in the 1970s to bypass damaged or blocked fallopian tubes and circumvent some cases of infertility, research into the female reproductive tract has tended to neglect the oviduct. As a result, our knowledge of the epithelial cell types within the oviduct, their developmental origins and homeostatic mechanisms remain poorly understood. Recently, interest into oviduct epithelial cells has been reignited by the discovery that epithelial cells from the distal tip of the fallopian tube are in many cases, the cell-of-origin in high-grade serous ovarian carcinoma (HGSOC) patients, the most common and deadly form of ovarian cancer (Kroeger & Drapkin, 2017; Labidi-Galy et al., 2017; Y. Lee et al., 2007; Medeiros et al., 2006; Shaw, Rouzbahman, Pizer, & Pintilie, 2009; Soong et al., 2018). Characteristics of the cell-of-origin plays a key role in determining cellular pliancy and subsequent pathophysiology of developing tumours (Choi et al., 2012; Hoadley et al., 2018; Puisieux et al., 2018; Zhu et al., 2016). In addition, the process of malignant transformation can take decades until cancer cells acquire uncontrolled proliferation and metastatic ability (Gerstung et al., 2020). This is tightly linked with changes in tissue homeostasis to accommodate changes in the genomic landscape of a cell. A fundamental understanding of oviduct epithelial heterogeneity and homeostasis is therefore required in order to investigate the early development of HGSOC and develop accurate models of the disease.

Morphologically the oviduct is divided into four evolutionary conserved regions: most distally are the infundibulum and ampulla, located adjacent to the ovary followed by the isthmus and uterotubal junction (Agduhr, 1927). Lining the oviduct lumen is an epithelium of secretory (PAX8+) and multiciliated (FOXJ1+) cells. The proportion of multiciliated cells is highest in the infundibulum and ampulla, decreasing along the length of the oviduct (Stewart & Behringer, 2012). Lineage tracing experiments of *Pax8* expressing cells in mice has shown that these cells are proliferative and can give rise to more PAX8+ secretory and multiciliated cells, suggesting PAX8+ cells are bipotent progenitors (Ghosh et al., 2017). It is currently unknown if there is further diversification of the epithelial cell types within the morphologically distinct regions of the oviduct.

There is evidence of functional differences between the distal (infundibulum and ampulla) and proximal (isthmus and uterotubal junction) regions of the oviduct. In an *ex vivo* study into the binding of sperm to the oviduct epithelium, distinct mechanisms of sperm binding to multiciliated epithelial cells was found in cultured segments of the ampulla and isthmus (Ardon et al., 2016). In addition, an *in vitro* study into the role of the oviduct in oocyte maturation showed that co-cultures of oocytes with epithelial cells isolated from the ampulla significantly increased zona pellucida hardening compared to those cultured with epithelial cells isolated from the isthmus (Dadashpour et al., 2015). Two transcriptional analyses of isolated epithelial cells from the ampulla and isthmus regions of the bovine oviduct also revealed extensive differences in gene expression between the two regions during the estrus cycle and pregnancy (Cerny et al., 2015; Maillo et al., 2010). In both studies however, bulk RNA sequencing of isolated epithelial cells was used, therefore it is unknown if the differential expression reported reflects different populations of secretory and multiciliated cells or changes in the proportion of these two cell types.

Developmentally the oviduct is derived from the anterior portion of the Müllerian duct which gives rise to the oviduct, uterus, cervix and anterior portion of the vagina (Yesmin et al., 2018). In mice, the Müllerian duct forms around E11.5 (Embryonic day) with the invagination of specified coelomic epithelial cells (Orvis & Behringer, 2007). The Müllerian duct then continues to extend rostro-caudally to the urogenital sinus adjacent to and dependent on signals released by the Wolffian duct (Atsuta & Takahashi, 2016; Kobayashi, Shawlot, Kania, & Behringer, 2004; Mullen & Behringer, 2014). Patterning of the Müllerian duct into the different structures of the female reproductive tract is then directed postnatally by spatially restricted expression of *Hox* genes. *Hoxa9* specifying oviduct development in mice (Benson et al., 1996; Satokata et al., 1995; Taylor et al., 1997; Warot et al., 1997). The mechanism determining further subcompartmentization of the oviduct into the morphologically distinguishable infundibulum, ampulla, isthmus and uterotubal junction is still unknown.

In the present study we identified heterogeneity in secretory and multiciliated cells and spatially mapped these subpopulations to different locations along the mouse oviduct. We identified differential expression of genes important for fertilization and embryonic development, and distinct expression patterns of classical cell type specific markers including PAX8, PAX2 and WT1 between proximal and distal regions. Distal and proximal populations formed early in Müllerian duct development, prior to previously identified postnatal restricted of *Hox* gene expression (Taylor et al., 1997). In addition, using lineage tracing and organoid culture of isolated epithelial cells we determined that distal and proximal epithelial cells are intrinsically different lineages and maintained separately. Taken together these results demonstrate that oviductal epithelial cells located in the distal and proximal regions are developmentally distinct lineages reflecting their unique roles and homeostasis in reproduction and potential susceptibility to tumorigenesis.

## Results

### *Pax2* expression distinguishes the distal and proximal regions of the mouse oviduct

*Pax2* expression is an early marker of specified Müllerian duct cells and is required for urogenital development (Torres et al., 1995). PAX2 and PAX8 are generally considered as secretory cell specific transcription factors in the adult oviduct. In order to investigate the expression pattern of *Pax2* in the mouse oviduct we used a *Pax2-GFP BAC* transgenic mouse line in which GFP expression faithfully reports *Pax2* expression (Figure 1A) (Pfeffer et al., 2002). PAX2-GFP expression was observed in the proximal region of the oviduct but surprisingly decreased at the ampulla-isthmus junction and was completely absent from the distal oviduct (Figure 1B). Transverse sections of the oviduct confirmed uniform expression of PAX2-GFP in all epithelial cells of the uterotubal junction and isthmus, a subpopulation of epithelial cells at the ampulla-isthmus junction and no PAX2-GFP expressing cells in the distal ampulla and infundibulum (Figure 1C). In the isthmus both multiciliated cells and secretory cells showed PAX2-GFP expression (Figure 1D). Multiciliated cells in the isthmus were also PAX8 positive, in clear contrast to the infundibulum and ampulla where PAX8 expression is restricted to non-multiciliated cells (Figure 1D). The transition from PAX8-to PAX8+ multiciliated cells occurred at the ampulla-isthmus junction, coinciding with the appearance of PAX2-GFP expressing cells (Supplementary figure 1A). To validate the reliability of the *Pax2-GFP BAC* transgenic, we confirmed differential expression of *Pax2* by RT-PCR between the distal and proximal regions of the oviduct and immunostaining using a PAX2 specific antibody (Supplementary figure 1B-D). Taken together these observations showed there are at least four populations of epithelial cells in the mouse oviduct: Distal PAX2-/Pax8-multiciliated cells, distal PAX2-/Pax8+ non-multiciliated cells, proximal PAX2+/Pax8+ multiciliated cells and proximal PAX2+/Pax8+ non-multiciliated cells.

**Figure 1.**
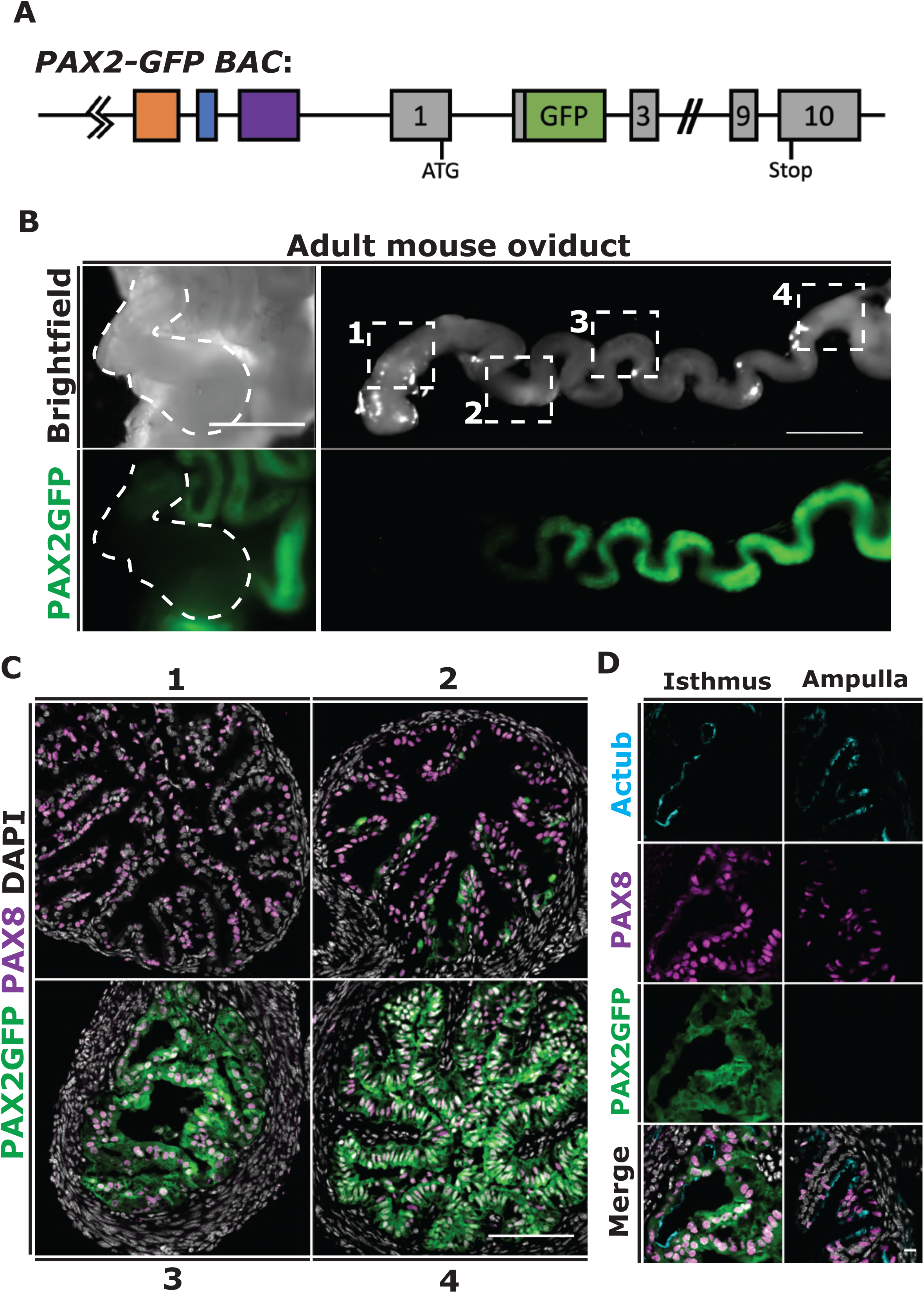
*Pax2* expression is restricted to the proximal region of the oviductal epithelium. *Pax2* expression in the mouse oviduct was investigated using a *Pax2-GFP Bac* transgenic mouse line (Pfeffer et al., 2002). **(A)** *Pax2-GFP Bac* mice have extra copies of the *Pax2* locus with an in-frame GFP fused to the end of exon2 and enhancer elements indicated by the coloured boxes 30kb upstream of the *Pax2* gene. **(B)** Whole mount imaging of an adult mouse oviduct coiled and uncoiled, showing absence of GFP expression in the distal region. **(C)** Transverse sections of indicated regions in **B** confirms localization of PAX2GFP expression in proximal epithelial cells. **(D)** High magnification of transverse sections showing expression of PAX2GFP and PAX8 in both secretory and multiciliated cells in the isthmus and secretory cell restricted PAX8 expression in the ampulla. Scale bars = 1000µm in B and 100µm in C and D.

### Single-cell RNA sequencing reveals heterogeneity in functional cell types

In order to investigate heterogeneity, we used single-cell RNA sequencing of oviduct epithelial cells isolated from an adult female in estrus (see methods). 807 epithelial cells were identified by expression of *Epcam* and clustered by T-distributed Stochastic Neighbor Embedding (t-SNE) into six populations (Supplementary Figure 2A-B). Clusters 0-4 and 1-2 showed a high degree of similarity in the top differentially expressed genes defining these populations. To confirm the independences of these populations we conducted a differential expression analysis comparing these populations directly. Clusters 0-4 had a total of 1028 differentially expressed genes (log2FC > 0.5, FDR < 0.05). However, clusters 1-2 showed no significantly differentially expressed genes (log2FC > 0.5, FDR < 0.05), which lead us to merge these clusters for the remainder of our analysis resulting in 5 clustered populations (Figure 2A). The expression of multiciliated cell marker *FoxJ1*, and secretory cell marker *Ovgp1*, broadly separated the clusters into two multiciliated and three secretory cell populations (Figure 2B and C). Highly differentially expressed genes were identified as defining each cluster (average log fold change > 0.5 and p < 0.001) (Figure 2D and Table 1). *Pax2* and *Pax8* were not detected at high levels in our scRNA-seq data (Supplementary figure 2C and D), likely due to the high dropout rates seen in shallow single-cell sequencing reflecting relatively low expression levels of these transcription factors (Chen et al., 2019).

**Figure 2.**
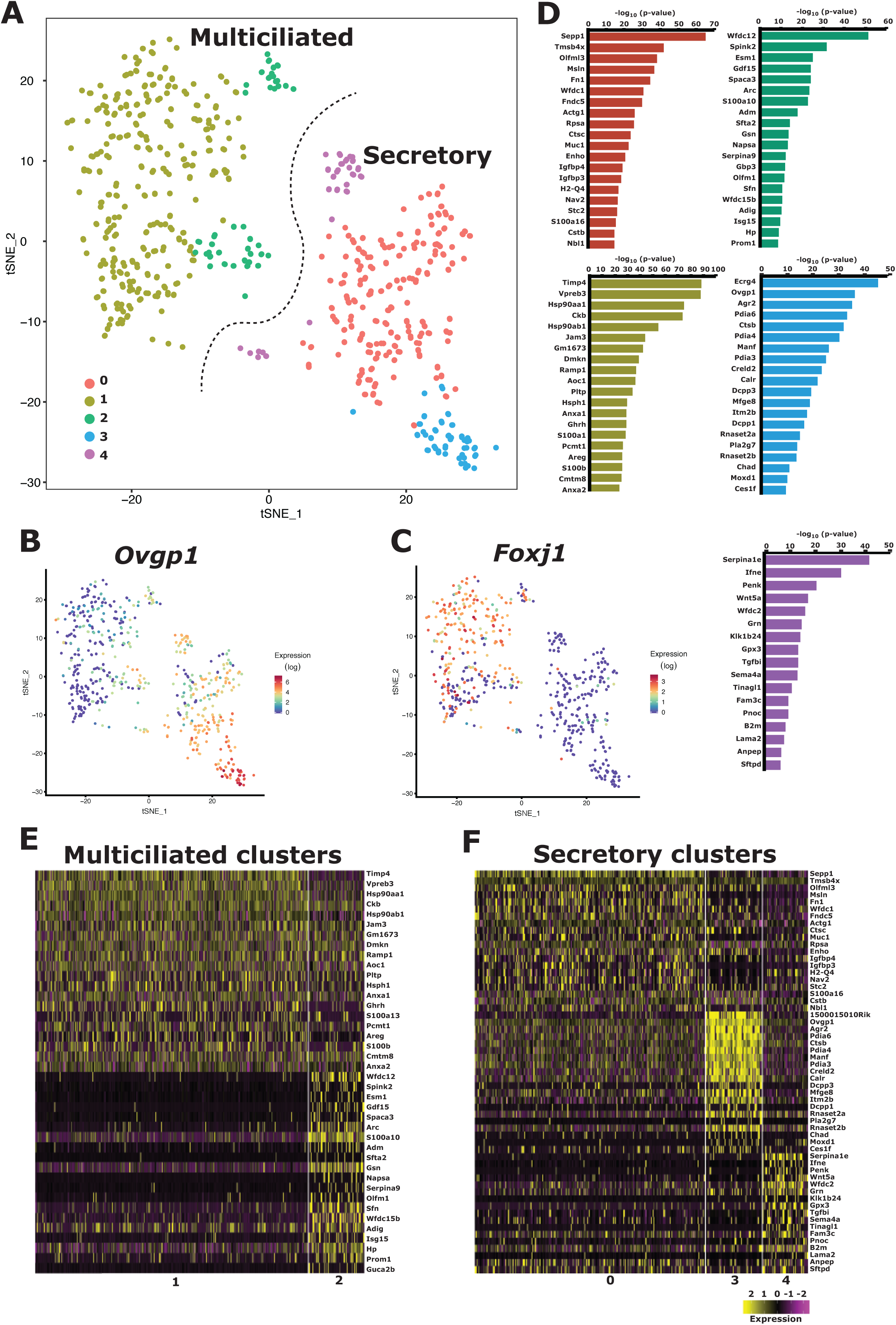
Single-cell RNA sequencing identifies five distinct epithelial cell populations. Mouse oviduct epithelial cell heterogeneity was investigated using single-cell RNA sequencing of epithelial cells isolated from a mouse in estrus. **(A)** 470 epithelial cells were identified and clustered using tSNE into 5 distinct clusters. **(B and C)** Heat maps showing the expression of multiciliated cell marker *Foxj1* and secretory cell marker *Ovgp1*. **(D)** Top 20 differentially expressed genes defining each cluster (log2FC > 0.5, FDR < 0.05) (See Table 1 for full list). **(E and F)** Heat maps comparing the top 20 differentially expressed “secreted” and/or “extra-cellar space” genes between multiciliated and secretory clusters.

In the oviduct, proteins important for reproduction are frequently secreted or bounded to the apical membrane of epithelial cells in order to support sperm, oocytes or developing embryos survival and migration. To investigate the functional differences between the epithelial populations during estrus, proteins with a GO term annotation of “secreted” and/or “extracellular space” were filtered using the Mouse Genome Informatics database (MGI) (Figure 2E and F, Table 2). We identified many significantly differentially expressed genes (average log fold change > 0.5 and p < 0.001) within each cluster including genes proposed to be involved in embryo development and oocyte maturation: *Sepp1, Igfbp3, Igfbp4* and *Pltp* (Burk et al., 2013; Lee et al., 2005; Wang et al., 2006). Sperm binding, survival and guidance: *Fn1, Hsp90aa1, Hsp90ab1, Anxa2, Nppc* and *Penk* (Almiñana et al., 2017; Ignotz et al., 2007; Kong et al., 2017; Osycka-salut et al., 2017; Subiran et al., 2012). Fertilization: *Ovgp1* and *Pdia6* (Avilés et al., 2010; Shuai Li & Winuthayanon, 2017). Protease inhibitors: *Wfdc1, Wfdc12, Wfdc15b, Wfdc2, Serpina9* and *Serpina1e* (Ranganathan et al., 2000; Winuthayanon et al., 2015) and antioxidant activities: *GPX3* and *Sod3* (Agarwal et al., 2014; Guerin et al., 2001; Lapointe & Bilodeau, 2003). Taken together these results indicate that there is functional heterogeneity in the secretory and multiciliated cell populations lining the mouse oviduct.

### Epithelial subpopulations are spatially restricted along the length of the mouse oviduct

To spatially map the epithelial populations identified in our scRNA-seq analysis we identified four markers whose differential expression could be used to define each cluster: *FoxJ1, Wt1, Ly6a/Sca1* and *Gpx3* (Figure 3A). We then immunostained for the selected markers along the length of the mouse oviduct during estrus to determine the location of each population. WT1 expression was identified in all distal epithelial cells in the infundibulum and ampulla but absent from the proximal epithelial cells in the isthmus and uterotubal junction in opposition to PAX2-GFP expression (Figure 3B). Based on the distal specific expression pattern of WT1 we were able to spatially map clusters 0 and 1 as PAX8+:WT1+ secretory and PAX8-:WT1+ multiciliated cells respectively, located in the distal oviduct (infundibulum and ampulla) (Figure 3C). Clusters 2 and 3 were mapped as PAX8+:WT1-secretory and PAX8+:WT1-:Ly6a+ multiciliated cells, located in the isthmus (Figure 3D). Cluster 4 was defined by high GPX3 expression and was mapped as a PAX8+:GPX3+ secretory cell population located in the uterotubal junction (Figure 3E). Gene expression in the oviduct is regulated by hormonal changes during the estrus cycle (Buhi et al., 2000). In order to determine if the specificity of our markers is maintained throughout the estrus cycle, we repeated the immunostaining analysis using oviducts from mice in diestrus. The specificity of the selected markers and location of cell populations for clusters 0-3 was maintained in diestrus (Supplementary figure 3A-D). However, GPX3 expression became less restricted in diestrus and was found in epithelial cells along the entire length of the oviduct (Supplementary figure 3E). The dynamic expression of GPX3 is in line with previous findings in bovine where high levels of GPX3 protein was found at the uterotubal junction during estrus, which progressively became more diffuse along the length of the oviduct through the estrus cycle (Lapointe & Bilodeau, 2003). Encouragingly, several cell type specific genes we identified by scRNA analysis have previously been shown to have regional specific expression in concordance with our results. *Nppc*, identified in distal secretory cells, has been shown to be highly expressed in the mouse ampulla and to act as an attractant for sperm migration (Kong et al., 2017). *Adm*, identified in multiciliated cells of the isthmus, has been shown to increase cilia beat frequency in the presence of sperm and is highly expressed in the human isthmus (H. W. R. Li et al., 2010).

**Figure 3.**
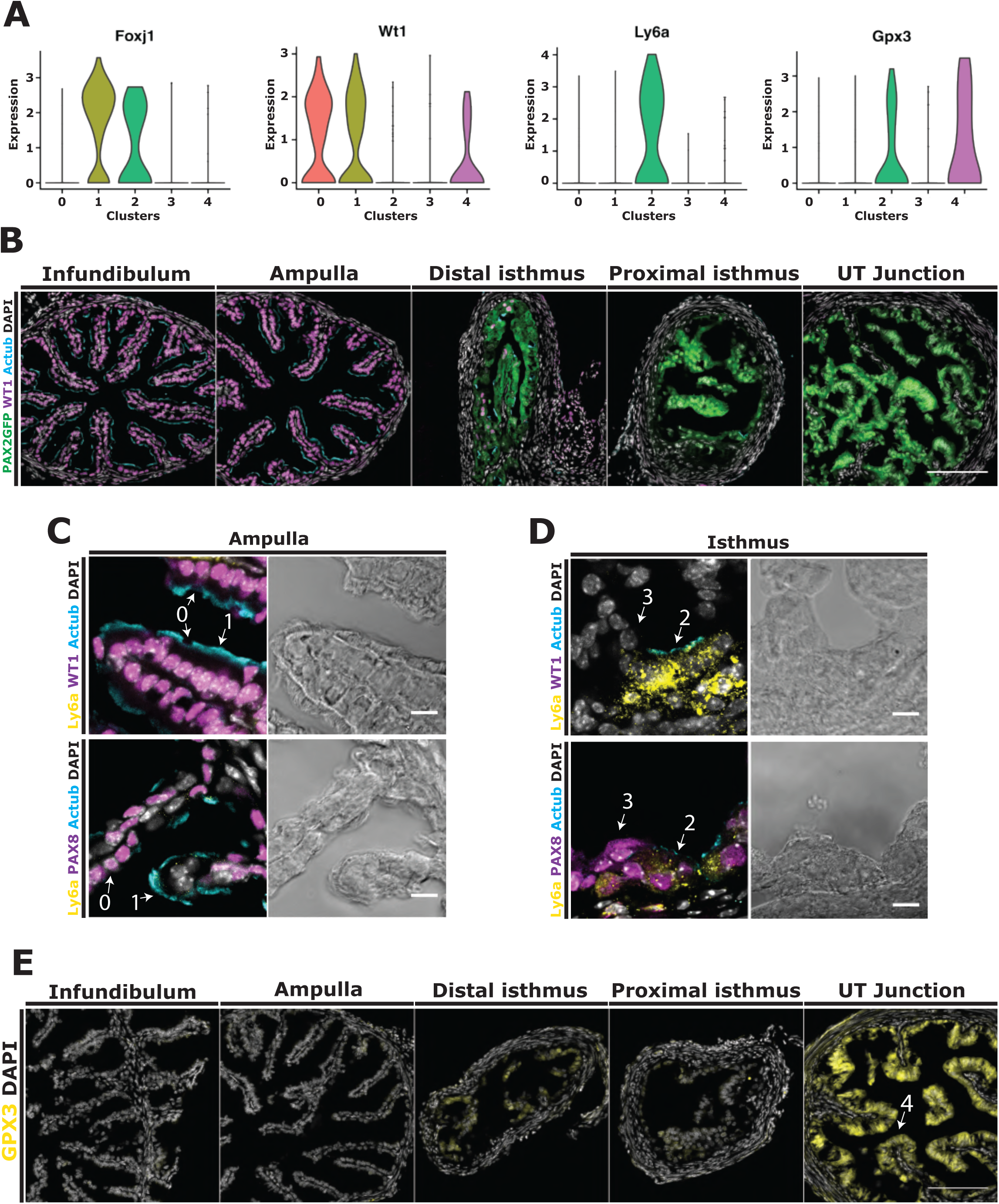
Spatially mapping oviduct epithelial cell populations. Epithelial cell populations were spatially mapped in the mouse oviduct during estrus using immunofluorescence. **(A)** Violin plots showing the expression of marker genes *FoxJ1, Wt1, Ly6a/Sca1* and *Gpx3* in each cluster. **(B)** Transverse sections taken along the length of the mouse oviduct showing expression of WT1 restricted to epithelial cells in the distal oviduct, opposing PAX2GFP expression. **(C)** High magnification image of the mouse ampulla indicating cluster 0 (secretory) cells (Wt1+:PAX8+:GPX3-) and cluster 1 (multiciliated) cells (WT1+:PAX8-:Ly6a-). **(D)** High magnification image of the mouse isthmus indicating cluster 2 (multiciliated) cells (WT1-:PAX8+:Ly6a+) and cluster 3 (secretory) cells (WT1-:PAX8+:GPX3-). **(E)** Transverse sections along the length of the oviduct showing high expression of GPX3, a markers of cluster 4 cells, in epithelial cells of the uterotubal junction. Scale bars in B and E = 100 µm, C and D = 10 µm.

### Distal-proximal specification occurs prior to epithelial cell differentiation in the mouse oviduct

To identify the origins of heterogeneity, we examined if distal-proximal specification occurred before epithelial differentiation. In the mouse, oviduct epithelial differentiation into mature cell types commences with the emergence of multiciliated cells at P4 (postnatal day 4) followed by secretory cell differentiation at P6 (Agduhr, 1927; Ghosh et al., 2017). We therefore undertook a second round of scRNA-seq using oviduct epithelial cells isolated from a single litter of P4 mice (4 mice, 8 oviducts) (see methods). 263 epithelial cells were identified by the expression of *Epcam* and clustered into two populations with significantly differentially expressed genes (Figure 4A and B, Table 3). *Wt1*, which we previously found to be specifically expressed in the distal adult oviduct epithelial cells (Figure 3A and Supplementary figure 3A), was identified to be one of the top differentially expressed genes between the two clusters (average log fold change = 0.44 and p <=0.02), suggesting clusters 0 and 1 may represent distal and proximal oviduct progenitors respectively. Distal restricted expression of *Wt*1 was confirmed in P4 oviducts and was found to be opposing to PAX2-GFP expression confirming distal-proximal specification occurs prior to epithelial differentiation (Figure 4C).

**Figure 4.**
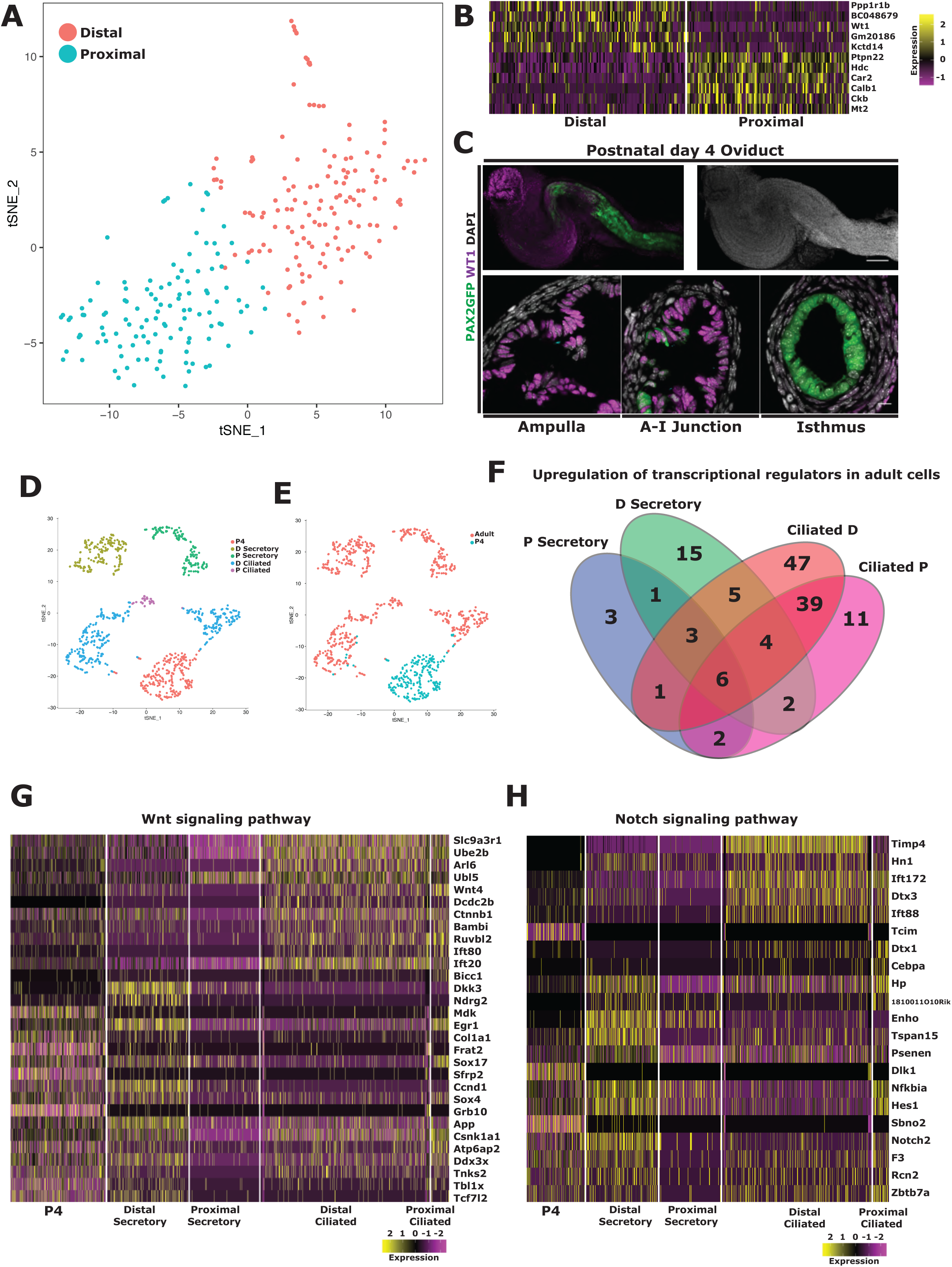
Distal and proximal cell identity is determined before oviduct epithelial differentiation. The presence of distal-proximal specification prior to epithelial differentiation was determined by scRNA sequencing of 263 epithelial cells isolated from P4 embryos. **(A)** TSNE plot of P4 epithelial cells showing clustering into two populations. **(B)** A heat map of the top differentially expressed genes identified *Wt1*, an adult distal cell marker, to be differentially expressed. **(C)** Whole-mount Z-projection and transverse sections confirm distal restricted expression of WT1 before epithelial cell differentiation. **(D)** To investigate gene expression changes during differentiation, adult and P4 datasets were combined and clustered by tSNE. **(E)** Some P4 cells clustered in the adult distal multiciliated cluster suggesting differentiation was starting to occur. **(F)** A Venn diagram showing the number of upregulated transcriptional regulators between P4 and adult populations. Heat maps showing the differentially expressed genes between P4 and adult populations found in the Wnt **(G)** and Notch **(H)** signaling pathways (MGI GO-term). Scale bar in C = 100 µm in top panels and 10 µm in bottom panels.

It has been shown in transgenic mice and organoid cultures that oviduct epithelial cell differentiation is controlled by a combination of Wnt and Notch signaling (Ghosh et al., 2017; Kessler et al., 2015; Xie et al., 2018). However, the precise genes involved in directing differentiation *in vivo* have not been identified. In order to find potential transcriptional regulators important for epithelial differentiation we combined our adult and P4 data sets to identify genes dynamically expressed during differentiation (see methods). TSNE clustering segregated the P4 and adult epithelial cell populations (Figure 4D and E). Distal adult multiciliated cell clusters were merged as before and cells from the uterotubal junction clustered with other proximal secretory cells. Several cells from the P4 sample clustered with adult distal multiciliated cells suggesting that some differentiation had occurred. We performed a differential expression analysis, independently comparing P4 cells to each of the differentiated clusters and filtered the results by GO terms relating to “transcription” (MGI) (Table 4). More transcriptional regulators were differentially expressed in the multiciliated populations compared to secretory populations, which were more similar to undifferentiated P4 cells (Figure 4F). In the multiciliated populations we identified many regulators previously identified to be essential for multiciliated cell differentiation including *Foxj1* and *Fank1* (Brody et al., 2000; Johnson et al., 2018). Several regulators were found to be unique to each population such as *Ghrh* and *Chek2*, and *Cebpb* and *Atf3* in distal and proximal populations respectively (Table 4). We also identified several members of the Wnt and Notch signaling pathways differentially expressed including: *Wnt4, Dkk3, Notch2* and *Dlk1* (Figure 4G and H).

### The distal oviduct epithelial population forms during Müllerian duct formation

To investigate the origins of distal-proximal populations, we traced the expression pattern of WT1 and PAX2, during Müllerian duct development. At E11.5, the onset of the Müllerian duct formation by invagination, we observed double PAX8 and PAX2-GFP expressing cells within the coelomic epithelium adjacent to the developing Wolffian duct (Figure 5A and Supplementary figure 4A). Consistent with previous reports, cells of the coelomic epithelium showed high WT1 expression (Armstrong et al., 1992). WT1 expression was however lower in cells expressing PAX2-GFP compared to surrounding cells (Figure 5A and B). By E12.5, WT1 expression was absent from the extending Müllerian duct, which was identified by PAX2-GFP expression adjacent to the Wolffian duct (Figure 5C and E). The cells at the rostral end of the Müllerian duct, that will form the distal end of the oviduct, were disorganised with intermingled WT1 and PAX2-GFP expressing cells, and a small population of WT1+:PAX2GFP-cells at the rostral tip (Figure 5D). The rostral compartment, marked by an absence of PAX2-GFP expression, continued to expand with Müllerian duct extension (Figure 5F and Supplemental figure 4B).

**Figure 5.**
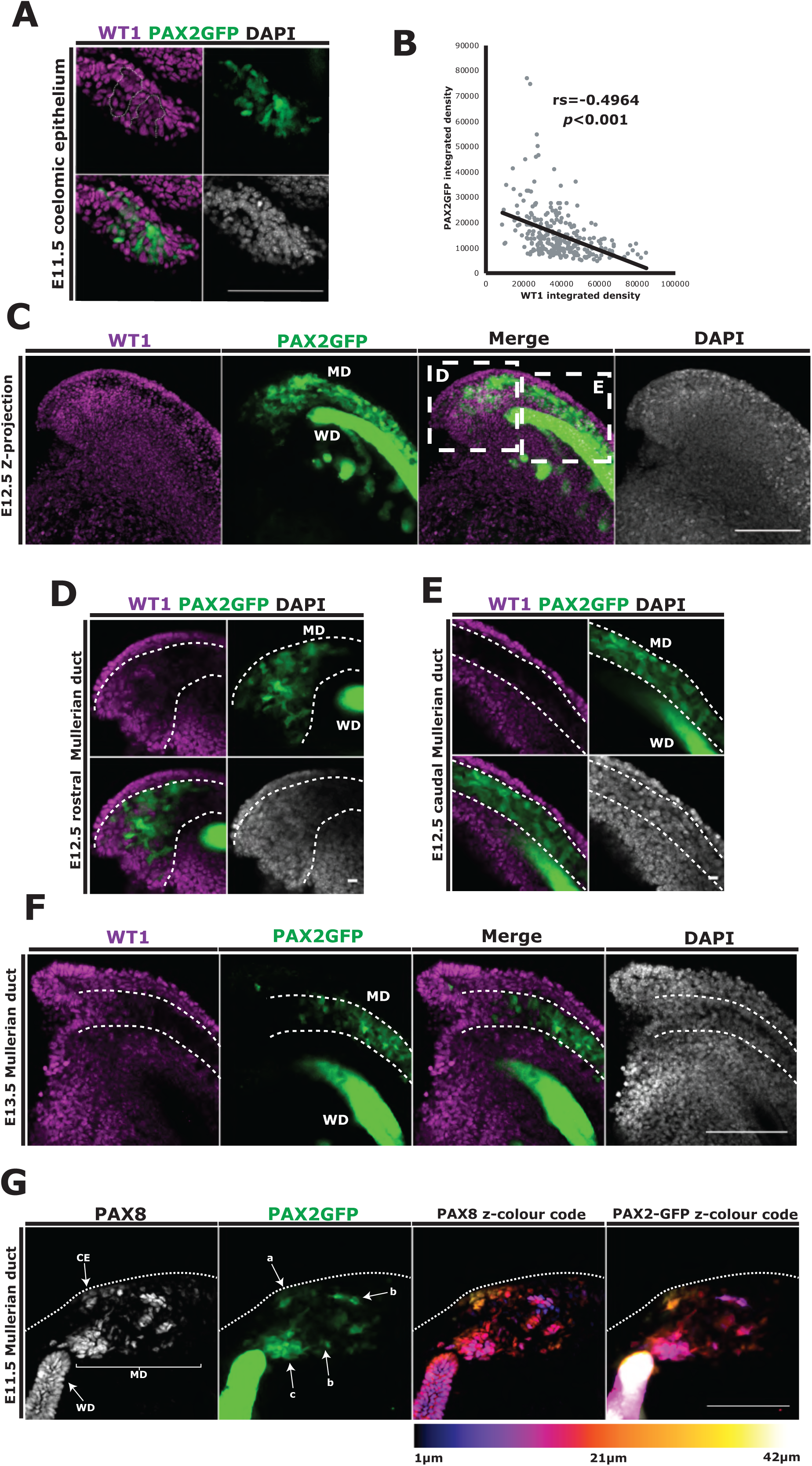
Distal oviduct epithelial cell population can be recognized in the Müllerian duct by E12.5. The formation of distal-proximal lineages was investigated by tracing the expression pattern of WT1 and PAX2 during Müllerian duct development in PAX2GFP mice. **(A)** Whole-mount image of the mesonephros at E11.5 showing specified Müllerian duct cells expressing PAX2GFP. **(B)** Specified cells in the coelomic epithelium showed a down regulation of WT1 expression compared to surrounding PAX2GFP negative cells. **(C)** Whole-mount z-projection of the mesonephros at E12.5 revealed the Müllerian duct lateral to the Wolffian, identifiable by PAX2GFP expressing epithelial cells. **(D and E)** A single z plane from the distal and proximal regions of the Müllerian duct highlighted in C. **(F)** A single z plane from the distal portion of the Müllerian duct at E12.5. A PAX2GFP-:WT1+ population is clearly visible at the rostral tip of the Müllerian duct. **(G)** A z-projection of the invaginating Müllerian duct on E11.5. Nuclear shape is highlighted by PAX8 staining and the cell body by cytoplasmic PAX2-GFP. Cells are then coloured by z-position to illustrate the 3D orientation of invaginating cells. Scale bars in A, C, F and G = 100 µm. Scale bars in D and E = 10 µm. Inverse relationship in C confirmed by 2-tailed Spearman’s Rank, rs = -0.4964 and *p*<0.001. n = 3 mice, 278 cells.

The cellular events of Müllerian duct initiation are poorly understood and have not been investigated in mice. In the chick, a sequential process of epithelial thickening, apical constriction and invagination has been identified (Atsuta & Takahashi, 2016). Contribution of epithelial to mesenchymal transition (EMT) is also suggested. To visualise the onset of Müllerian duct formation, we performed high resolution 3D confocal imaging of invaginating Müllerian ducts at E11.5 using PAX2GFP mice counter stained with a PAX8 antibody (Figure 5G and Supplementary figure 4C). PAX8+:PAX2GFP+ cells were identified in the coelomic epithelium (labelled “a” in Figure 5G and Supplementary figure 4C z = 18 µm). A group of cells with mesenchymal-like characteristics including elongated cell bodies, stretched nuclei and cytoplasmic projections stretched from the celomic epithelium through the stromal compartment of the mesonephros towards the Wolffian duct (labelled “b” in Figure 5G and Supplementary figure 4C z = 18 µm). A packed cluster of PAX8+: PAX2GFP+ cells within the stromal compartment showed higher PAX2GFP expression compared to surrounding cells formed adjacent to the Wolffian duct. (labelled “c” in Figure 5G and Supplementary figure 4C z = 12 µm). A small group of PAX2GFP+ cells extended caudally in the direction of future Müllerian duct elongation behind the Wolffian duct (labelled “d” in Supplementary figure 4C z = 42 µm). Our results suggest that initiation of the Müllerian duct is not simply regulated by invagination but also by EMT after cell specification in the coelomic epithelium.

### Distal/proximal oviductal epithelial cells are distinct lineages and separately maintained

In order to investigate the homeostasis of distal and proximal populations, we performed a lineage tracing study using a knock-in *PAX8*^*Cre*^ and *Rosa26*^*Tdt*^ Cre-reporter lines (Bouchard et al., 2004; Madisen et al., 2010). In the developing Müllerian duct PAX8 is uniformly expressed in epithelial cells (Supplementary figure 4B) and becomes restricted to secretory cells in the oviduct during epithelial differentiation around P4 (Ghosh et al., 2017). However, cre recombinase activity of the *PAX8*^*Cre/+*^ allele is reported as mosaic in some tissues due to low expression levels, that can be overcome by making the allele homozygous (Bouchard et al., 2004). On E14.5 in *PAX2GFP*:*PAX8*^*Cre/+*^:*Rosa26*^*Tdt*^ embryos, we identified mosaic expression in the Müllerian duct and a scattering of Tdtomato+ cells (Figure 6A). By E17.5 clusters of Tdtomato+ cells were seen in both distal and proximal regions of the oviduct, recognizable by differential PAX2GFP expression. Postnatally we continued to see a scattered pattern of Tdtomato+ cells until P3. Interestingly, from P4 we observed an enrichment of Tdtomato+ cells in the distal portion of the oviduct, complementary to PAX2GFP expression and coinciding with timing of the appearance of multiciliated cells (Figure 6A) (Agduhr, 1927; Ghosh et al., 2017). In agreement with a previous lineage tracing study using *Pax8-rtTA;tetO-Cre* mice (Ghosh et al., 2017), we observed Tdtomato expression in both secretory and multiciliated cells (Figure 6B). The enrichment of Tdtomato+ cells in the distal region, where no PAX2GFP expression was observed, as well as relatively low numbers of scattered Tdtomato+ cells in the proximal region, where PAX2GFP was expressed, were maintained throughout the life of the mouse (confirmed up to 13 months of age) (Figure 6C-E). The consistent distribution pattern of Tdtomato+ cells indicates that distal epithelial cells do not contribute to the posterior population. This also suggested that the distal and proximal populations follow distinct differentiation trajectories and are maintained as independent lineages. To validate that distal and proximal populations are distinct lineages, not affected by the surrounding stromal environment, we generated organoids from mouse oviduct epithelial cells using culture conditions previous described (Xie et al., 2018) (Figure 7A). After 7 days in culture, organoids could be seen from both regions (Figure 7B). Organoids derived from the distal region were larger (mean diameter 92.9±11.4µm from distal cells compared to 61.3±5.6µm from proximal cells), and grow at a higher frequency when seeded at identical densities, 31.9±4.9 cells per mm^3^ compared to 13.5±4.7 cells per mm^3^ (n = 5 mice) (Figure 7C and D). In addition, we found that regional specific markers PAX2 and WT1 for proximal and distal, respectively, were maintained in culture (Figure 7E and F). When differentiation was induced by addition of γ-secretase inhibitor, cilia were identified in organoids from both regions (Figure 7G and H). The regional identity of epithelial cells was maintained *in vitro* without support from the surrounding stromal environment. Taken together, distal and proximal oviduct epithelial cells are two distinct and separately maintained lineages.

**Figure 6.**
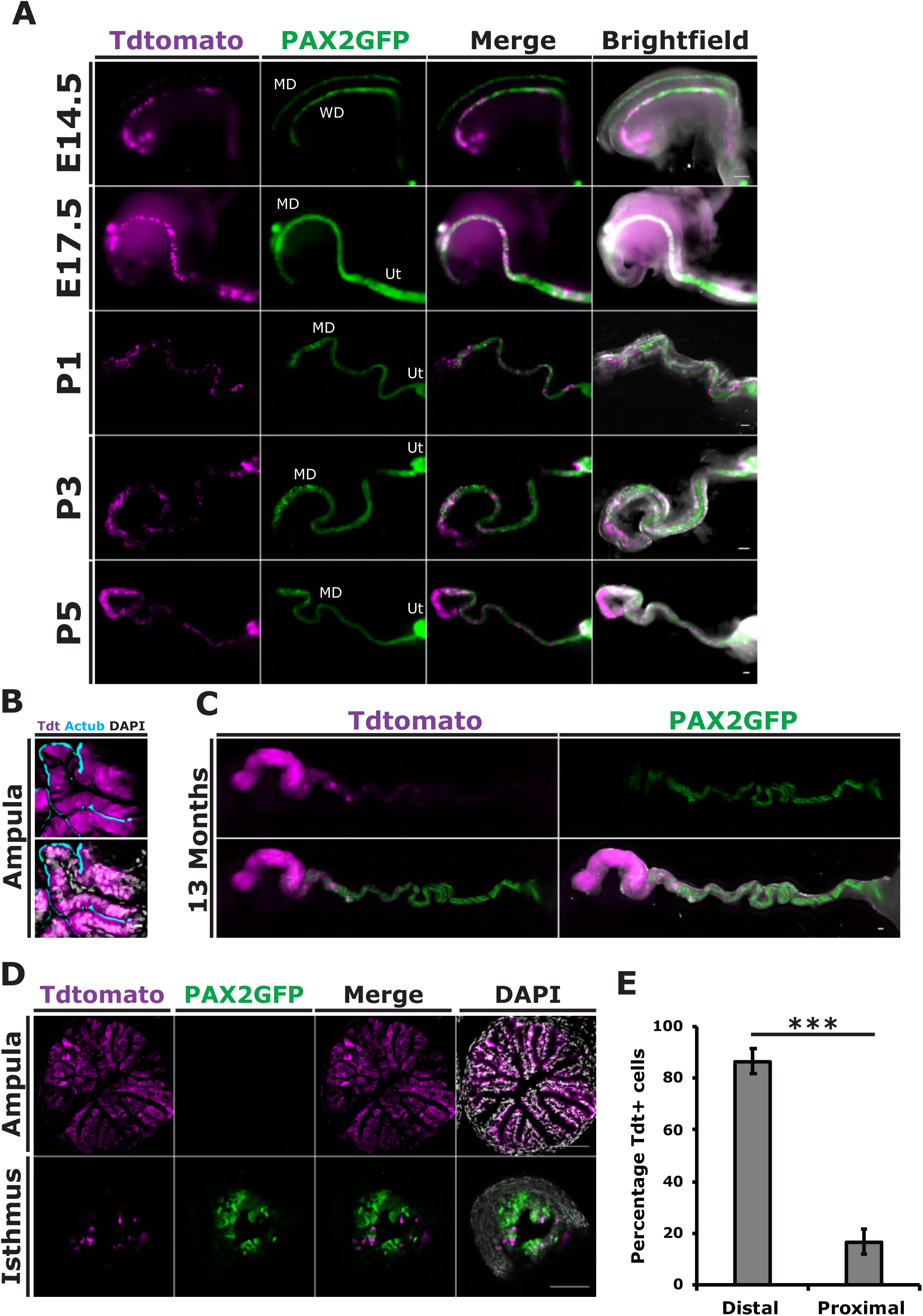
Distal and proximal oviduct epithelial cells are maintained separately. Oviduct epithelial homeostasis was investigated by mosaic lineage tracing of *Pax8* expressing cells using *PAX2GFP*:*PAX8*^*Cre/+*^:*Rosa26*^*Tdt/Tdt*^ mice. **(A)** Whole-mount images of the developing oviduct from E14.5 to P5. Enrichment of Tdtomato+ cells can be seen in the distal oviduct from P3. **(B)** Section of the adult ampulla showing Tdtomato+ multiciliated and secretory cells. **(C)** Distal enrichment of Tdtomato+ cells was maintained in adult mice, confirmed to 13 months. **(D)** Transverse section through the ampulla and isthmus in adult mice confirming the distal enrichment of Tdtomato+ cells. **(E)** Quantification of Tdtomato+ cells between distal and proximal regions. Significant difference in E confirmed by a 2-tailed student’s T-test, *P*<0.001, n = 10 mice. Scale bars = 100 µm in A,C,D and 10 µm in B.

**Figure 7.**
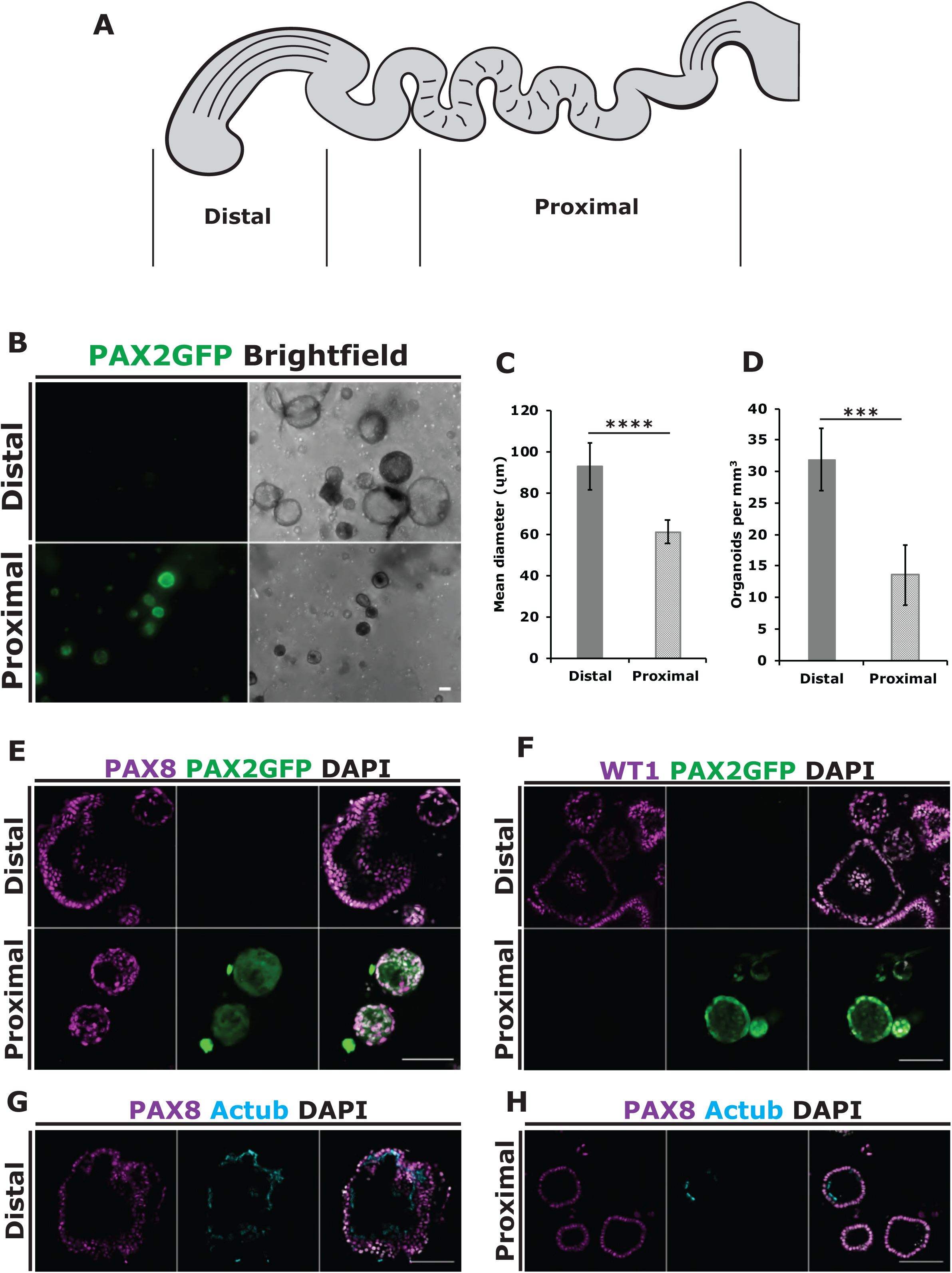
Mouse oviduct epithelial organoids maintain their regional identity. **(A)** Mouse oviduct organoids were generated from the distal and proximal regions of the oviduct, excluding cells from the ampulla isthmus junction. **(B)** Images of organoids derived from PAX2GFP females after 7 days in 3D culture. Expression of PAX2GFP was only observed in organoids grown from the proximal region. **(C)** Distal organoids were significantly larger than organoids isolated from the proximal region (n = 337 distal organoids and 143 proximal organoids from matched cultures). **(D)** Distal epithelial cells also developed into organoids at a higher frequency when seeded at the same density (n = 7 fields of view from matched cultures). **(E)** Confocal images of organoids showing expression of PAX8 in all epithelial cells and PAX2GFP specifically in all epithelial cells of proximal organoids. **(F)** Expression of distal specific marker WT1 was found only in organoids isolated from distal epithelial cells. **(G** and **H)** Multiciliated cell differentiation induced in organoids by addition of 1µM selective γ-secretase inhibitor. Multiciliated cells formed after 5 days in distal **(G)** and proximal organoids **(H)**. Scale bars = 100 µm.

## Discussion

The oviduct is essential for reproduction, providing an environment to promote fertilization and support preimplantation development (Shuai Li & Winuthayanon, 2017). These functions are primarily carried out by secretory and multiciliated epithelial cells lining the luminal surface of the oviduct. Multiciliated cells generate flow, important for the transport of gametes and embryos and have also been shown to physically bind with sperm in many species for sperm storage (Lyons et al., 2006). Secretory cells produce a mixture of proteins, nutrients and exosomes which interact with gametes and embryos as they traverse through the oviduct (Shuai Li & Winuthayanon, 2017; Maillo et al., 2016). It has long been considered that the epithelial cells within the oviduct are a single population of cells with two functional cell types (secretory and multi-ciliated). Using scRNA sequencing and analysis of markers PAX2/8 and WT1, we identified discrete populations of secretory and multiciliated cells restricted along the distal-proximal axis of the oviduct (Figure 8A). These populations had differential expression of many proteins with known functions during reproduction and provides evidence of the functional differences between the distal and proximal regions of the oviduct across species (Ardon et al., 2016; Cerny et al., 2015; Dadashpour et al., 2015; Maillo et al., 2010). Importantly, distal and proximal oviductal epithelial cells are specified early in the Müllerian duct and maintained separately. In addition, they have an intrinsic cellular identity independent from the surrounding stromal cells. Taken together, the oviductal epithelium consists of two distinct lineages, distal and proximal, both having the potential to differentiate into multiciliated and secretory cells.

**Figure 8.**
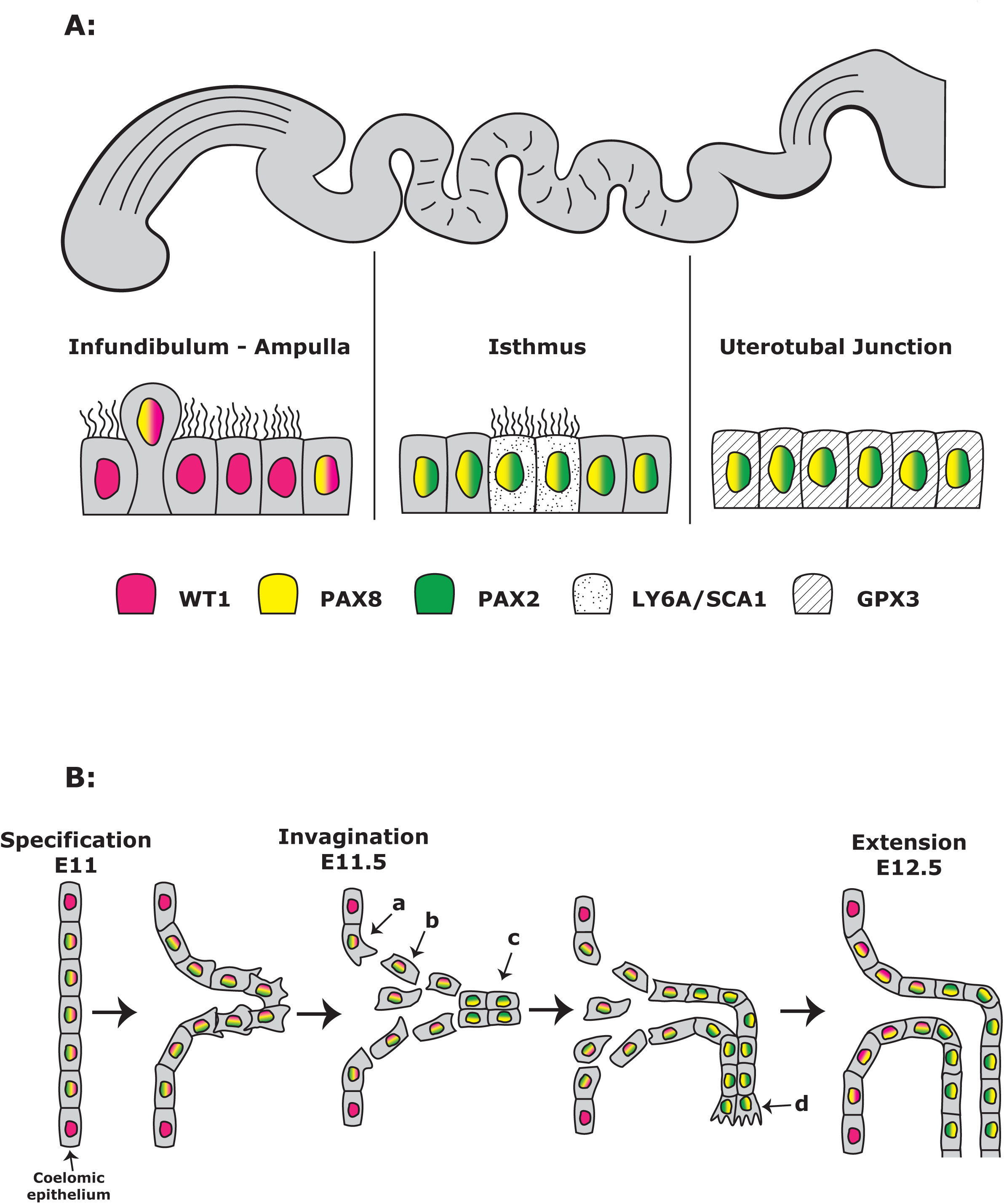
Models of oviduct epithelial cellular heterogeneity and its development. **(A)** The mouse oviduct is morphologically segregated into infundibulum, ampulla, isthmus and uterotubal junction. Epithelial cells within the distal (infundibulum and ampulla) and proximal (isthmus and uterotubal junction) are maintained separately as distinct lineages, expressing different markers and functional genes. **(B)** The two distinct epithelial lineages form early in Müllerian duct development. At E11 coelomic epithelial cells are specified as precursor cells for the Müllerian duct, upregulating PAX2/8 while downregulating WT1. Müllerian duct precursor cells at E11.5 undergo a process of invagination and EMT/MET. Müllerian duct precursor cells form a condensed cluster of cells close to the Wolffian duct which then extend caudally. By E12.5 WT1 expression is lost in the majority of Müllerian duct cells but for a population of cells at the rostral end, which becomes the distal end of the oviduct, maintaining WT1 but no PAX2 expression. This region expands with Müllerian duct extension.

In mice, it has been thought that coelomic epithelial cells expressing *Lim1* between E11-E11.5 invaginate to form the Müllerian duct (Kobayashi et al., 2004). However, our 3D confocal microscopy analysis at the onset of Müllerian duct initiation identified signs of epithelial mesenchymal transition (EMT). At E11.5, we identified four PAX2GFP+ populations marking precursors of the Müllerian duct: a) GFP+ cells within the coelomic epithelium, b) GFP+ mesenchymal-like cells within the stromal compartment of the mesonephros, c) a packed cluster of high GFP+ cells adjacent to the Wolffian duct and d) a small group of GFP+ cells towards the direction of extension (Figure 8B indicated as a,b,c and d respectively). During gastrulation, epiblast cells invaginate and then migrate out from epiblast to form mesoendoderm by EMT (Nakaya & Sheng, 2008). Similar to gastrulation, the precursor cells forming the Müllerian duct are specified on the surface coelomic epithelium. Our results suggest a process of EMT/MET could lead to the formation of a packed aggregate/rod underneath the coelomic epithelium along the Wolffian duct, which will elongate similar to nephric duct development (Grote et al., 2006; K. Stewart & Bouchard, 2014). Both WT1 and PAX2 have been implicated in EMT/MET in development and are thought to repress each other in kidney development (Doberstein et al., 2011; Hastie, 2017; B. Huang et al., 2012; Martínez-estrada et al., 2010; Rothenpieler & Dressler, 1993). It is interesting if these transcriptional factors are also involved in similar processes during Müllerian duct formation, in addition to their functions in cell specification.

During invagination, we found WT1 expression is down regulated in PAX2GFP+ cells resulting in the formation of a WT1 negative, PAX2:PAX8 positive cell population. As the Müllerian duct continues to elongate around E12.5, a distinct population, WT1:PAX8 positive, PAX2 negative, was recognized at the rostral end of the elongating Müllerian duct. This is prior to sexual specification and *Hox* gene expression is still homogeneous (Taylor et al., 1997). The formation of distinct lineages at this time may reflect independent origins of distal-proximal populations. Alternatively, early and late specified coelomic epithelium cells may progress through different cellular states during tube formation imparting distal-proximal characteristics.

In the adult oviduct our analysis identified two types of multiciliated and secretory cells that are restricted to distal and proximal regions. Differential expression of genes with known functions in modulating sperm behaviour including: *Itm2b, Penk, Nppc* and *Fn* was found between secretory cell populations (H. Choi et al., 2015; Kong et al., 2017; Osycka-salut et al., 2017; Subiran et al., 2012). Indicating compartmentalised protein secretions that modulate sperm behaviour along the length of the oviduct. In a parallel study (Harwalkar et al., n.d.), we identified unique structures created by multiciliated cells in the isthmus. Multiciliated cells are clustered in the trench of transverse luminal folds forming pits or grooves, clearly distinct from the uniform distribution of multiciliated cells on the longitudinal luminal folds in the infundibulum and ampulla. Consistent with these unique structures, we identified differentially expressed genes involved in sperm binding, *Anxa1, Anxa2* and *Adm*, suggesting distinct sperm-cilia interactions in the isthmus (Ardon et al., 2016; Ignotz et al., 2007; H. W. R. Li et al., 2010). Further functional studies comparing distal and proximal epithelial cells are required to identify their specific roles during reproduction.

Serous Tubal Intraepithelial Carcinoma (STIC) lesions in the distal oviduct have been identified as the origin of HGSOC in familial and 50% of sporadic cases (Kroeger & Drapkin, 2017; Labidi-Galy et al., 2017; Y. Lee et al., 2007; Soong et al., 2018). It has been proposed that the formation of STIC lesions limited in the distal oviduct is a result of repeated exposure to reactive oxygen species and cytokines released by the ovary (H. Huang et al., 2015; Jones et al., 2013). Alternatively, there have been several reports showing the accumulation of progenitor cells at the distal region of the oviduct that could be predisposed to malignant transformation (Alwosaibai et al., 2017; Ng et al., 2013; Paik et al., 2012; Patterson & Pru, 2013; Y. Wang et al., 2012; Xie et al., 2018). Our study demonstrates that the distal epithelial cells are a distinct lineage from the rest of the oviduct. It is interesting to speculate that the distal oviductal epithelium could be uniquely susceptible to oncogenic insults. A recent study also identified heterogeneity in human fallopian tube epithelial cells and was able to link the signature of a subtype of secretory cells to poor prognoses in HGSOC patients (Hu et al., 2020). It will be important to consider the cell-of-origin and its impact on tumor development/pathophysiology for further understanding of ovarian cancer heterogeneity.

Our findings lay a foundation for understanding morphological regionalities and their distinct functions within the oviduct. Further research will be required to determine factors essential for distal and proximal lineages and how they respond to female hormonal cycles in addition to their contribution in fertilization and preimplantation development.

## Acknowledgements

We thank the McGill Goodman Cancer Research Centre Histology, Flow Cytometry facilities and the McGill Advanced Bioimaging Facility (ABIF) and Comparative Medicine and Animal Research Centre (CMARC). We also thank Genome Quebec for sequencing. This work was supported by Canadian Cancer Society (CCS) Innovation grant (Haladner Memorial Foundation #704793), CCS i2I grant (# 706320) and Cancer Research Society Operation Grant (#23237). M.J.F was supported by Canderel, CRRD and FRQS postdoc fellowships. K.H. was supported by CRRD and Alexander McFee (Faculty of Medicine) and Gosselin studentships. K.T. was supported by MICRTP and Canderel studentships. J.R is supported by Genome Canada Genome Technology Platform grants and the Canada Foundation for Innovation.

## Author contributions

Y.Y conceived the study. M.J.F performed the majority of the experiments and prepared the manuscript. Y.Y, K.H and M.J.F edited the manuscript. K.H, H.M and K.T performed some immune stainings and imaging. A.S.P analyzed the single cell RNA sequencing data. Y.C.W prepared single cell RNA sequencing libraries for sequencing and performed QC analysis. D.B aided with analysis of single cell data. J.R supervised scRNA sequencing experiments, QC, parts of the data analysis and contributed to project design. N.Y maintained mouse colonies. M.B provided *PAX2-GFP* and *PAX8Cre* mice and shared unpublished information.

## Methods

### Mouse stains and maintenance

All animal work was approved by the internal ethics committee at McGill University and undertaken at the Goodman Cancer Research Centre animal facility. *Pax2-GFP Bac* and *Pax8Cre* transgenic mice (Bouchard et al., 2004; Pfeffer et al., 2002) were received as a kind gift from Prof. Maxime Bouchard. *Tdtomato*^flox/flox^ mice (Ai14) were acquired from JAX (#007914). C57BL/6 stock mice were used as wild type mice for scRNA sequencing and immunofluorescent stainings. Adult mice were analysed during reproductive age and the estrus cycle staged by vaginal smear followed by crystal violet staining (Mclean et al., 2012).

### Immunofluorescence

Embryonic Müllerian and adult oviducts were dissected in PBS and fixed in 4% w/v PFA/PBS (Polysciences) for 30 minutes at room temperature followed by 3 PBS washes. Ducts were cryoprotected through a sucrose/PBS gradient 4°C, allowing time for the ducts to sink to the bottom of the tube between each gradient. Ducts were embedded in OCT mounting solution (Fisher HealthCare) and snap frozen on dry ice before being stored in a -80°C freezer. 10µm sections were cut using the microtome cryostat Microm HM525 (Thermofisher) mounted on Superfrost glass slides, air dried and stored in -80°C freezer. For immunofluorescence, sections were allowed to thaw for 10mins at room temperature followed by rehydration with PBS. Sections were permeabilized for 5 minutes with 0.5% v/v Triton-X/PBS (Sigma) and then blocked with 10% v/v Fetal Bovine Serum (FBS) (Wisen Bioproducts), 0.1% v/v Triton-X in PBS for 1 hour at room temperature. Primary antibodies were diluted 1/250 in 1% v/v FBS, 0.1% v/v Triton-X in PBS and incubated overnight in a dark humidified chambered at 4°C. The following antibodies were used in this study: PAX2 (Invitrogen #71-6000), PAX8 (proteintech #10336-1-AP), WT1 (Adcam #ab89901), LY6A/SCA1 (Abcam #ab51317), Acetylated Tubulin (Sigma #T7451) and GPX3 (R&D Systems (#AF4199). The followed day sections were washed before incubation with secondary antibodies ((Alexa Fluor) diluted 1/400 in 1% v/v FBS, 0.1% v/v Triton-X in PBS) for 1 hour in a dark humidified chambered at room temperature. Sections were then washed again with 5µg/ml DAPI (Fisher Scientific) added to the final wash step before sections were mounted with prolong gold (Invitrogen) for imaging. For whole mount immunofluorescence of embryonic Müllerian and postnatal oviducts, fixed samples were permeabilized for 10 minutes in 0.5% v/v Triton-X/PBS, stained as above and then mounted in prolong gold between two coverslips separated by a spacer (Invitrogen Sercure-seal 0.12mm).

### Reverse transcription-PCR analysis of *Pax2* expression

In order to semi-quantitatively compare the level of *Pax2* expression between the distal and proximal regions of the oviduct, the oviducts from adult mice were dissected into distal (infundibulum and ampulla) and proximal regions (isthmus and uterotubal junction) and RNA isolated using the RNA purification mini kit (Geneaid) following manufactures instructions with an on-column DNase I digest. Three mice (six oviducts) were combined into each RNA preparation. 500ng of RNA from each sample was then used to create cDNA using the iScript reverse transcription kit (Bio-rad) following manufactures instructions. PCR reactions for *Pax2* and *Gapdh* were set up using the 2X Taq FroggaMix (FroggaBio) with the following primers and performed using a Mastercycler ep gradient S (Eppendorf). *Pax2*: forward primer - GTACTACGAGACTGGCAGCATC, reverse primer – CGTTTCCTCTTCTCACCGTTGG (Beauchemin et al., 2011). *Gapdh*: forward primer - TGAGGCCGGTGCTGAGTATGTCG, reverse primer - CCACAGTCTTCTGGGTGGCAGTG (Shih-wein Li et al., 2016). A 6X loading dye (New England Biolabs) was added to each sample and the entire sample run on a 1% agarose gel with TAE buffer then imaged using a UV light and camera. Quantification of band intensity was performed using FIJI image analysis software.

### Isolation of mouse oviduct epithelial cells by FACS

Single cell suspensions of C57BL/6 P4 and adult oviduct epithelial cells were isolated by Fluorescence-activated cell sorting (FACS). Oviducts from a single litter of P4 mice (8 oviducts) and two oviducts from a single 3-month old mouse in estrus were disassociated into single cells by incubation in collagenase B (5mg/ml) and DNase I (5U/100ul) in DMEM (Gibco) containing 100 IU/ml of Penicillin and 100ug/ml of Streptomycin for 35 minutes at 37°C followed by passing through a needle series. The resulting single cell suspensions were passed through a 40µm cell strainer, centrifuged in a table top centrifuge at 1,500rpm for 5 minutes and resuspended in 200µl staining solution containing conjugated antibodies Ep-CAM-APC (#118213 Biolegend) and Brilliant violet 785-CD45 (#103149 Biolegend) diluted 1/150 with DMEM (Gibco) containing 100 IU/ml of Penicillin and 100ug/ml of Streptomycin and 1% FBS for 15 minutes in the dark on ice. Cell suspensions were then centrifuged, washed in PBS and resuspended in FACS media (DMEM (Gibco) containing 100 IU/ml of Penicillin and 100ug/ml of Streptomycin and 20% FBS). FACS was performed with the FACSAria Fusion (BD Biosciences) using FACSDIVA (Version 8). Cell suspensions were spiked with 1µl 5mg/µl DAPI or LIVE/DEAD stain (Thermofisher) just prior to sorting to identify dead or dying cells. Live Epcam+/CD45-epithelial cells were sorted into FACS media and concentrated for single cell cDNA library preparation by centrifugation.

### Quality control for scRNA-seq

An aliquot was taken from each sample cell suspension and incubated in live-dead staining consisting of working concentration of 2µM calcein-AM and 4µM Ethidium-Homodimer1 (Thermofisher L3224). After 10 minutes of incubation at room temperature, sample viability, concentration, segregation, size and absence of large debris were verified by loading stained cell suspension onto hemocytometer (Incyto DHC-N01-5) and imaged on bright field, GFP (for Calcein-AM) and RFP (for Ethidium homodimer-1) channels using a EVOS FL Auto Fluorescent microscope (ThermoFisher). Percent Viability is derived from GFP positive cells (live cells) over the sum of GFP positive plus RFP positive cells (all cells) and all samples with a viability bellow 70% were excluded.

### Generation of single cell cDNA libraries and sequencing

The single cell gene expression data was generated according to Chromium Single Cell 3’ Reagent Kits User Guide (v2 Chemistry, 10X genomics). Briefly, samples that passed quality control were suspended into Reverse Transcription (RT) Master Mix (10X genomics) then pipetted into Well-1 of a Chip “A” (10X genomics), followed by Gel Beads (10X genomics) into Well-2 and Partition oil (10X genomics) into Well-3. The chip assembly was run on a Chromium Controller (10X genomics) which generated Gel Bead-In-EMulsions (GEMs). GEMs were pipetted out from the Chip and into a 200µL PCR tube (Eppendorf 951010022) and ran on a thermocycler (Biorad T100) with RT protocol [45min at 53°C, 5min at 85°C, hold at 4°C]. After RT, the GEMs content was released using Recovery Reagent (10X genomics) and cDNA were isolated using Buffer Sample Clean Up 1 (10X genomics) containing Silane Dynabeads (Thermofisher 2000048). The purified cDNA was PCR amplified followed by purification using 0.6X volume SPRIselect beads (Beckman Coulter B23318). The cDNA quality (size distribution) and concentration were assessed using the LabChip (Perkin Elmer 760517, CLS760672). A one-step Fragmentation, End Repair & A-tailing mix (10X genomics) were added to the cDNA in a 200µL PCR tube and ran on a thermocycler with the protocol (hold at 4°C, 5min at 32°C, 30min at 65°C, hold at 4°C). The fragmented cDNA was subjected to double sided size selection using SPRIselect bead, by first suspending the fragmented cDNA in 0.6X volume of SPRIselect for 5 min, using 10x Magnetic Separator to pull down the beads, moving the suspension into another PCR tube and topped with 0.8X volume of SPRIselect. Following two rounds of 80% ethanol washes, the desired sized fragmented cDNA were eluted into adaptor ligation mix (10X genomics) and incubated for 15min at 20°C. The ligated product was cleaned with 0.8X volume of SPRIselect, added to Sample Index PCR Mix (10X genomics) and amplified via 12-14 PCR cycles depending on the amount of full-length cDNA as input (45sec at 98°C, (12 cycles of 20sec at 98°C, 30sec at 54°C, 20sec at 72°C), 1min at 72°C, hold at 4°C). The final PCR product (or sequence ready library) was purified just like fragmented cDNA and quality controlled using LabChip as described earlier. Finally, the libraries were sequenced on Illumina Hiseq4000.

### scRNA sequencing analysis

Raw sequencing data for each sample was converted to matrices of expression counts using the Cell Ranger software provided by 10X genomics (version 2.0.2). Briefly, raw BCL files from the Illumina HiSeq were demultiplexed into paired-end, gzip-compressed FASTQ files for each channel using Cell Ranger’s mkfastq. Using Cell Ranger’s count, reads were aligned to the mouse reference transcriptome (mm10), and transcript counts quantified for each annotated gene within every cell. The resulting UMI count matrix (genes × cells) were then provided as input to Seurat suite (version 2.3.4). Cells were first filtered to remove those that contain less than 200 genes detected and those in which > 10% of the transcript counts were derived from mitochondrial-encoded genes. Mean UMIs and genes detected in each cell were 3696 and 1546 for adult and 5499 and 2253 for P4 datasets. Clustering was performed using the “FindClusters” function and integrated t-SNE visualization for all cells. Cluster-specific gene markers were identified using Seurat’s FindMarkers with cutoffs avg_logFC > 0.5 and FDR < 0.05 unless otherwise stated. For the combined analysis, OVA or P4 samples were merged into one 10x combined object using canonical correlation analysis (CCA), followed by scaling data (ScaleData function) and finding variable genes (FindVariableGenes function). CCA subspaces were aligned using CCA dimensions 1-20, which was followed by clustering (FindClusters function) and integrated t-SNE visualization for all cells. Cluster-specific gene markers were identified using Seurat’s FindMarkers with cutoffs avg_logFC > 0.5 and FDR < 0.05. GO-term filtering was conducted using the Mouse Genome Informatic (MGI) database.

### Generation and culturing of mouse oviduct organoids

Mouse oviduct epithelial cells were isolated by first dissecting distal and proximal regions, as indicated in Figure 7A, in dissection solution (PBS containing 2% fetal bovine serum (FBS) and 100 IU/ml of Penicillin and 100ug/ml of Streptomycin). Oviducts were then broken into 1mm pieces with forceps before being incubated in collagenase B (5mg/ml) and DNase I (5U/100µl) in dissection solution for 35 minutes at 37°C. Oviducts were then spun for 5 mins at 250 G, resuspended in 100µl typsin and incubated for 5 minutes at 37°C. Oviducts were again spun for 5 mins at 250 G and resuspended in 200µl dissection solution. The resulting oviducts and cell suspensions were passed through a 18G needle 10 times to release epithelial cells and then through a 40µm cell strainer. Collected cells were resuspended in 100µl 2D culture media (DMEM/Ham’s F12 (Gibco #11320033), 15 mM HEPES (Gibco #15630080), 100 U/ml penicillin (Gibco #15140122), 100 µg/ml streptomycin (Gibco #15140122), 1X GlutaMAX (Gibco, #35050–061), 5% FBS (Wisent Bioproducts #090-150), 1X insulin-transferrin-selenium (Gibco #41400045), 25 ng/ml hEGF (Peprotech #AF-100-15-500), 30 µg/ml bovine pituitary extract (Gibco #13028014), and 20,000 cells plated per well of an uncoated 96 well plate. Epithelial cells were grown in a 37oC humidified incubator with 5% CO2 until 70% confluent (2-5 days). Epithelial cells were then seeded at a density of 25,000 cells per 50µl drop of Geltrex (Gibco #A1413201) in 24 well uncoated dishes which were allowed to set inverted for 30 minutes in a 37oC humidified incubator with 5% CO2 before being submerged in organoid media (DMEM/F12 advance media (Gibco #LS12634028) with HEPES (Gibco #15630080), 1X GlutaMAX (1:100, Gibco, #35050–061), 1X B27 (Gibco, #17504044), 0.1 µg/ml hEGF (PeproTech, #AF-100-15-500), 0.5µM TGFBR1 Kinase Inhibitor IV (SB431542, BioVision, #1674-1), 100 ng/ml mFGF10 (Peprotech, #450-61-25), 100 ng/ml hNoggin (StemCell Technologies, #78060.1), 0.6 µg/ml hR-spondin1 (StemCell Technologies, #78213.1) and 100 ng/ml mWNT3A (Peprotech, #315-20). For the first few days of culture after seeding and passaging 10 µM of Y-27632 (ROCK inhibitor, Sigma Aldrich, #Y0503) was added.

Organoids were passaged every 7-14 days and differentiation induced by the addition of 1 µM selective γ-secretase inhibitor Y0-01027 (DBZ) (Calbiochem, #209984-56-5) to the organoid culture media.

## Supplementary figures

**Supplementary figure 1.**
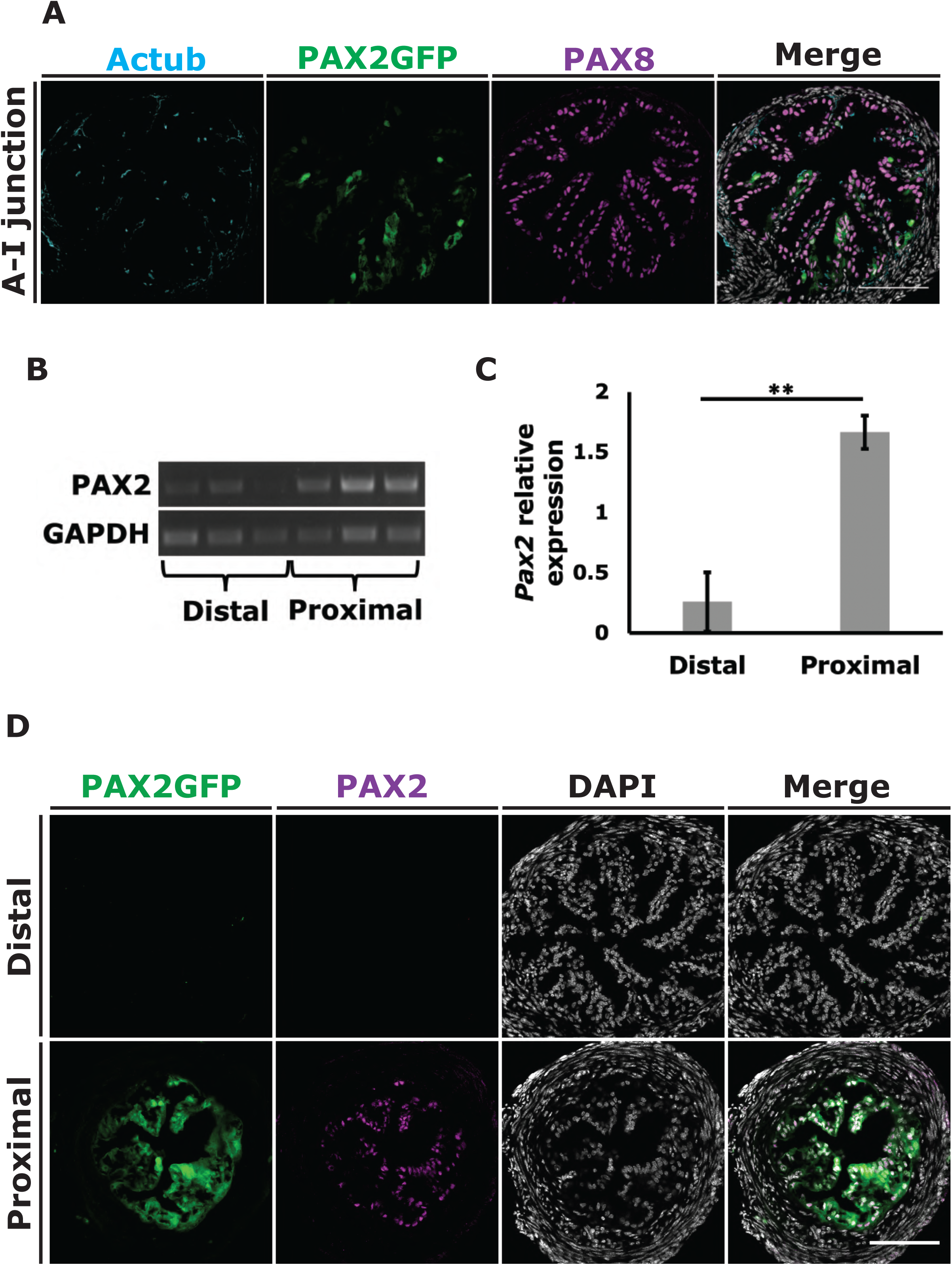
*Pax2-GFP Bac* expression faithfully reports *Pax2* expression in the mouse oviduct. PAX2GFP expression was identified specifically in the proximal oviduct with a boundary at the ampulla-isthmus junction **(A)** A transverse section at the ampulla-isthmus junction showing the transition to ubiquitous PAX8 specific expression in both secretory cells and multiciliated cells. The reliability of the PAX2GFP Bac to report *Pax2* expression in the oviduct was confirmed using RT-PCR and immunofluorescence. **(B)** RT-PCR comparing *Pax2* expression between distal and proximal regions of the oviduct (3 samples, 3 mice per sample). **(C)** Relative *Pax2* expression was calculated using expression of GPDH to normalise *Pax2* expression levels. *Pax2* expression was 6.641-fold higher in RNA isolated from the proximal region compared to the distal oviduct. **(D)** Immunofluorescence using a PAX2 antibody confirmed the absence of the PAX2 protein in the distal oviduct, corresponding with GFP expression. Significant difference in C confirmed by 2-tail Student’s T-test, *P*<0.005 Scale bar in C = 100 µm.

**Supplementary figure 2.**
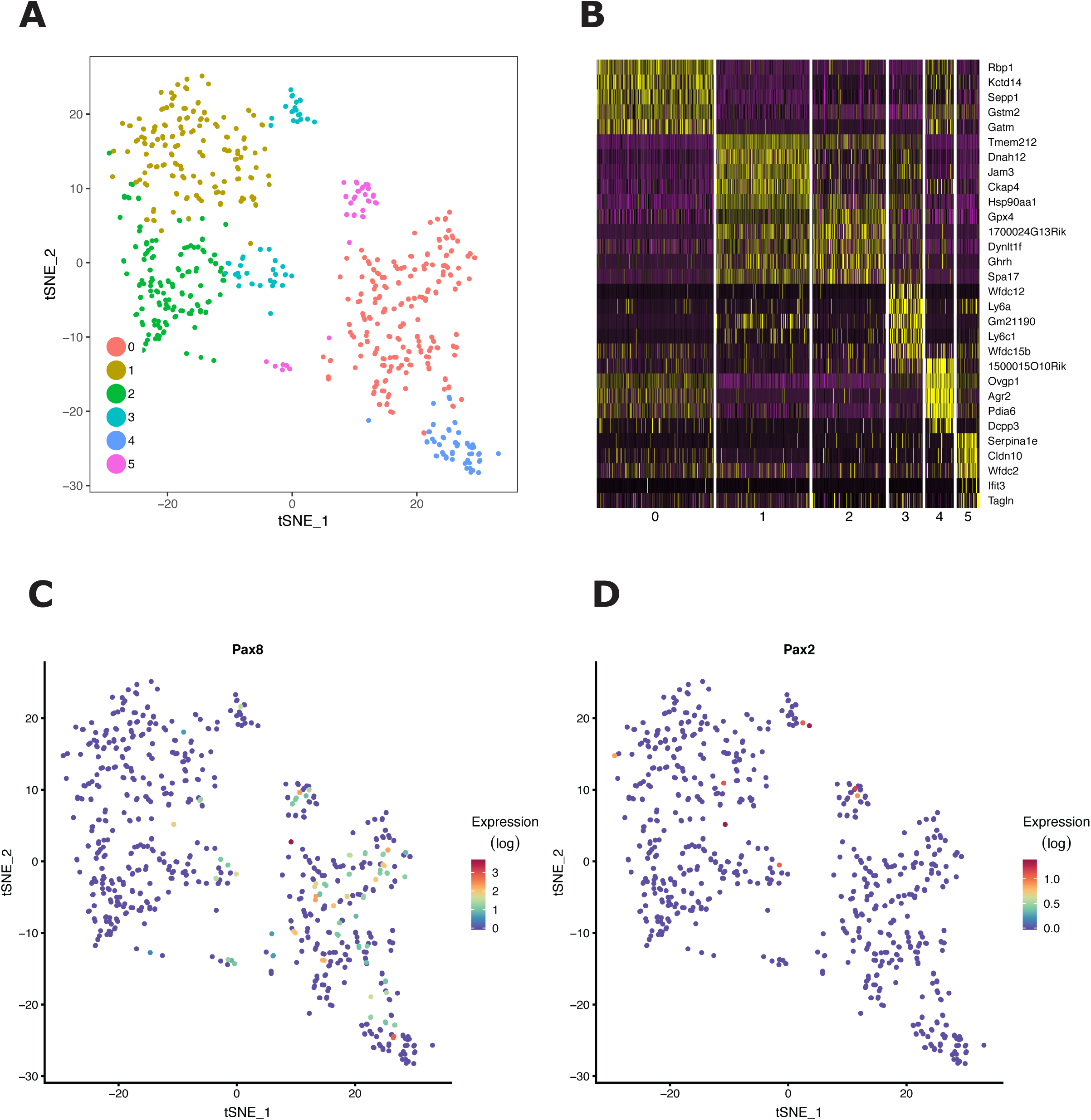
Clustering oviduct epithelial cells. Oviduct epithelial heterogeneity was identified using scRNA-seq. **(A)** Epcam+ epithelial cells were identified and clustered into 6 clusters using tSNE clustering. **(B)** A heat-map of the top differentially expressed genes defining each cluster in A, showing similar gene expression between clusters 1>2 and 0>4. **(C and D)** Due to shallow sequencing, markers *Pax8* and *Pax2* were not detected at high enough levels for further analysis.

**Supplementary figure 3.**
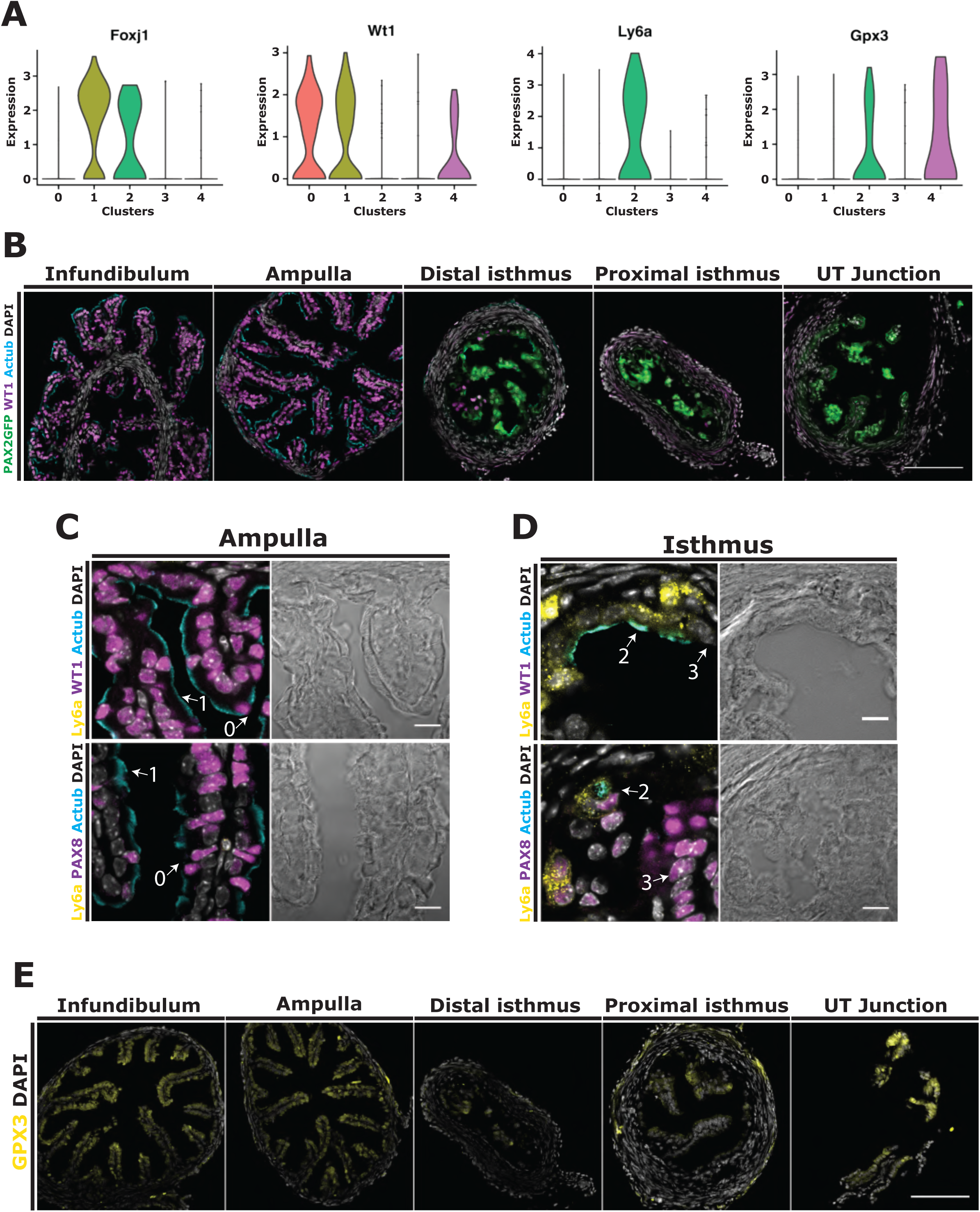
Distal and proximal markers are maintained in diestrus. The continuation of population markers in difference stages of the estrus cycle was investigated by immune-fluorescence staining oviducts isolated from mice in diestrus. **(A)** Violin plots showing the expression of markers *Foxj, Wt1, Ly6a* and *Gpx3* from the scRNA-seq of adult oviduct epithelial cells in estrus. **(B)** Transverse sections along the length of the mouse oviduct showing opposing WT1 and PAX2GFP expression, similar to the pattern seen in mice at estrus. **(C)** High magnification images of the ampulla showing the presence of WT1+:PAX8+ secretory and WT1+:PAX8-:Ly6a-multiciliated cells. **(D)** High magnification images of the isthmus showing WT1-:PAX8+ secretory and WT1-:PAX8+:Ly6a+ multiciliated cells. **(E)** GPX3 expression was identified in epithelial cells within all regions of the oviduct. Scale bars in B and E = 100 µm, C and D = 10 µm.

**Supplementary figure 4.**
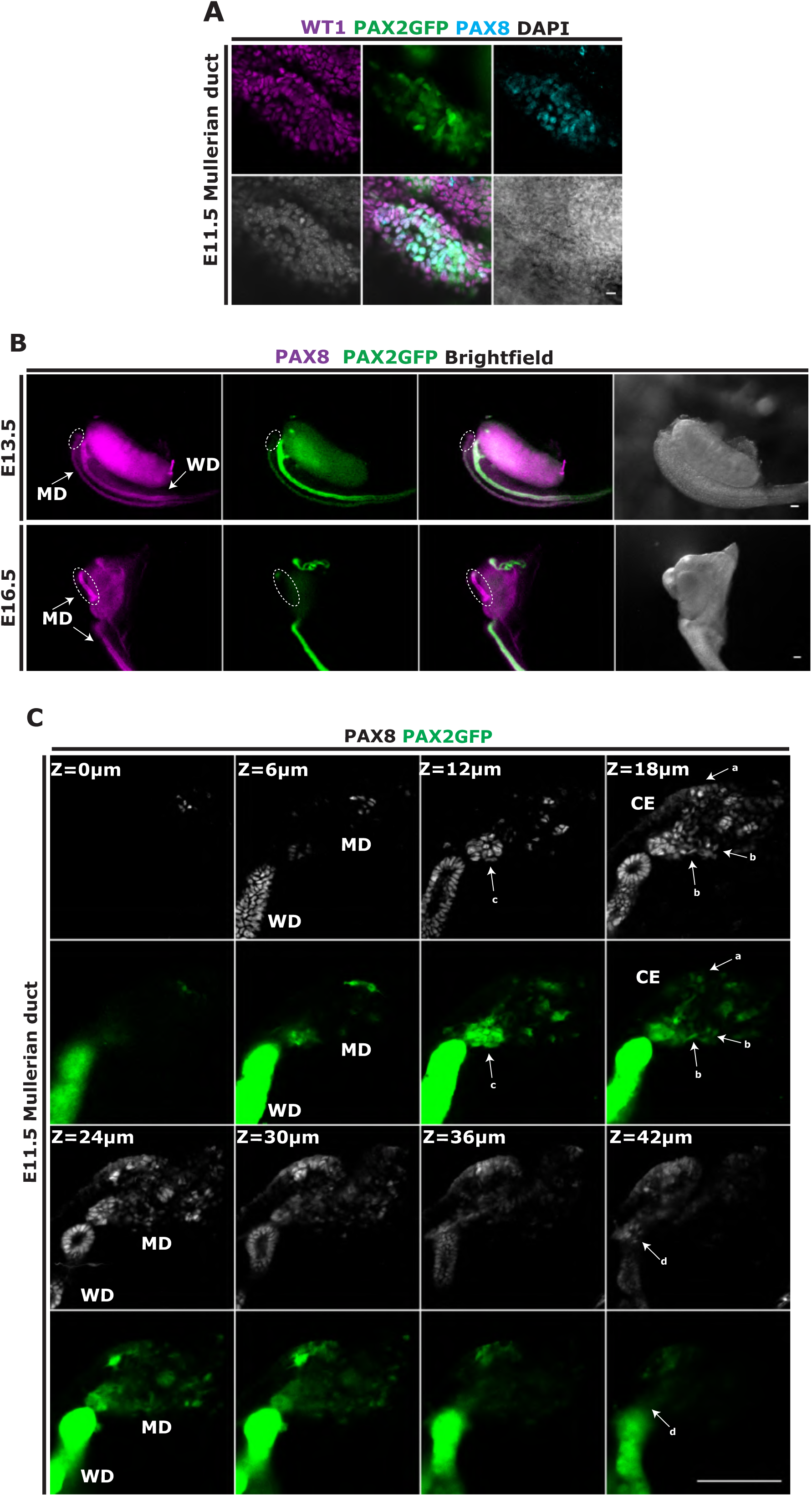
Initiation of Müllerian duct formation from coelomic epithelium. The behaviour of invaginating Müllerian duct precursor cells was investigated using whole-mount imaging. **(A)** An image of the mesonephros at E11.5 showing PAX8:WT1:PAX2-GFP expressing cells in the coelomic epithelium. **(B)** Images of the developing Müllerian duct on E13.5 and E16.5 showing expansion of the distal PAX2-GFP negative compartment. **(C)** Optical z sections passing through a whole-mount mesonephros showing the invaginating Müllerian duct precursor cells adjacent to the Wolffian duct on E11.5. (a) PAX8:PAX2GFP expressing cells on the coelomic epithelium. (b) A group of mesenchymal-like PAX8:PAX2GFP expressing cells. (c) A cluster of high PAX2-GFP expressing cells in close proximity to the Wolffian duct. (d) A group of PAX8:PAX2GFP expressing cells behind the Wolffian duct projecting caudally. Scale bars in A = 10 µm, B = 100 µm. MD = Müllerian duct, WD = Wolffian duct, CE = coelomic epithelium.

